# Unlinked rRNA genes are widespread among Bacteria and Archaea

**DOI:** 10.1101/705046

**Authors:** Tess E. Brewer, Mads Albertsen, Arwyn Edwards, Rasmus H. Kirkegaard, Eduardo P. C. Rocha, Noah Fierer

**Affiliations:** Cooperative Institute for Research in Environmental Sciences, University of Colorado, Boulder, CO 80309 USA; Department of Chemistry and Bioscience, Aalborg University, 9220 Aalborg, Denmark; Institute of Biological, Environmental and Rural Sciences, Aberystwyth University SY23 3DA UK; Microbial Evolutionary Genomics, Institut Pasteur, CNRS, UMR3525, Paris, 75015, France; Department of Ecology and Evolutionary Biology, University of Colorado, Boulder, CO 80309 USA

## Abstract

Ribosomes are essential to cellular life and the genes for their RNA components are the most conserved and transcribed genes in Bacteria and Archaea. These ribosomal rRNA genes are typically organized into a single operon, an arrangement that is thought to facilitate gene regulation. In reality, some Bacteria and Archaea do not share this canonical rRNA arrangement-their 16S and 23S rRNA genes are not co-located, but are instead separated across the genome and referred to as “unlinked”. This rearrangement has previously been treated as a rare exception or a byproduct of genome degradation in obligate intracellular bacteria. Here, we leverage complete genome and long-read metagenomic data to show that unlinked 16S and 23S rRNA genes are much more common than previously thought. Unlinked rRNA genes occur in many phyla, most significantly within Deinococcus-Thermus, Chloroflexi, Planctomycetes, and Euryarchaeota, and occur in differential frequencies across natural environments. We found that up to 41% of the taxa in soil, including dominant taxa, had unlinked rRNA genes, in contrast to the human gut, where all sequenced rRNA genes were linked. The frequency of unlinked rRNA genes may reflect meaningful life history traits, as they tend to be associated with a mix of slow-growing free-living species and obligatory intracellular species. Unlinked rRNA genes are also associated with changes in RNA metabolism, notably the loss of RNaseIII. We propose that unlinked rRNA genes may confer selective advantages in some environments, though the specific nature of these advantages remains undetermined and worthy of further investigation.

## Introduction

Ribosomes are the archetypal “essential proteins”, so much so that they are a key criteria in the division between cellular and viral life (Raoult and Forterre, 2008). In Bacteria and Archaea, the genes encoding the RNA components of the ribosome are traditionally arranged in a single operon in the order 16S - 23S - 5S. The rRNA operon is transcribed into a single RNA precursor called the pre-rRNA 30S, which is separated and processed by a number of RNases (Srivastava and Schlessinger, 1990). This arrangement of rRNA genes within a single operon is thought to allow rapid responses to changing growth conditions - the production of rRNA under a single promoter allows consistent regulation and conservation of stoichiometry between all three, essential components (Condon et al., 1995). Indeed, the production of rRNA is the rate-limiting step of ribosome synthesis (Gourse et al., 1996), and fast-growing Bacteria and Archaea accelerate ribosome synthesis by encoding multiple rRNA operons (Klappenbach et al., 2000).

Some Bacteria and Archaea have “unlinked” rRNA genes, where the 16S and 23S rRNA genes are separated by large swaths of genomic space (Figure 1). This unlinked rRNA gene arrangement was first discovered in the thermophilic bacterium *Thermus thermophilus* (Hartmann et al., 1987). Reports of unlinked rRNA genes soon followed in additional Bacteria, including the planctomycete *Pirellula marina* (Liesack and Stackebrandt, 1989), the aphid endosymbiont *Buchnera aphidicola* (Munson et al., 1993), and the intracellular pathogen *Rickettsia prowazekii* (Andersson et al., 1995). Though unlinked rRNA genes were first discovered in a free-living environmental bacteria, their ubiquity among the order Rickettsiales has led to suggestions that unlinked rRNA genes are a result of the genome degradation typical of obligate intracellular lifestyles (Rurangirwa et al., 2002; Merhej et al., 2009; Andersson and Andersson, 1999).

**Figure 1:**
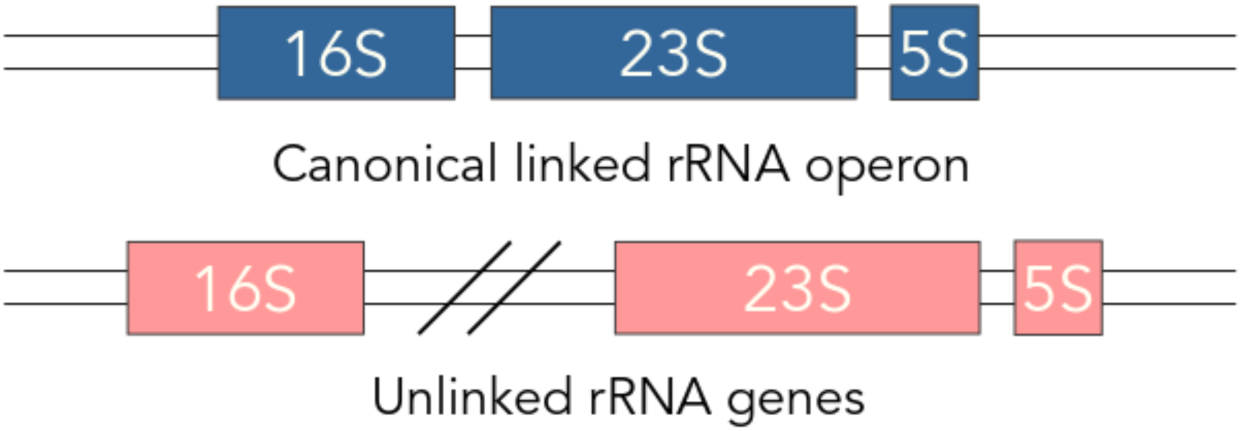
In most Bacteria and Archaea, the rRNA genes are arranged in the order 16S - 23S - 5S, and are transcribed and regulated as a single unit. However, in some cases, the 16S is separated from the 23S and 5S, and is referred to as “unlinked”.

With this study, we sought to determine the frequency of unlinked rRNA genes across Bacteria and Archaea and whether this unique genomic feature is largely confined to those Bacteria and Archaea with an obligate intracellular lifestyle. We examined the rRNA genes of over 10,000 publicly available complete bacterial and archaeal genomes to identify which taxa have unlinked rRNA genes and to determine if there are any genomic characteristics shared across taxa with this feature. As complete genomes are not typically available for the broader diversity of Bacteria and Archaea found in environmental samples (Zhi et al., 2012), we also characterized rRNA gene arrangements using long-read metagenomic datasets obtained from a range of environmental samples, which together encompassed over 17 million sequences (≥1000bp). With these long-read metagenomic datasets, we were able to determine whether unlinked rRNA genes are common in environmental populations and how the distributions of unlinked rRNA genes differ across prokaryotic lineages and across distinct microbial habitats.

## Methods

### Analyses of complete genomes

We downloaded all bacterial and archaeal genomes in the RefSeq genome database (O’Leary et al., 2016) classified with the assembly level “Complete Genome” from NCBI in January 2019 (12539 genomes). We removed genomes from consideration that had non-numeric gene ranges (96 genomes), >20 reported rRNA genes (2 genomes), or an unequal number of 16S and 23S rRNA genes (219 genomes). This left us with a set of 12222 genomes. We used gene ranges associated with each open reading frame (ORF) to pair the 16S and 23S rRNA genes that were closest to each other in each genome. We then checked for gene directionality (sense/antisense) and calculated the distance between each pair, taking directionality into account (see Supplemental Figure S1 for more detail and a visual representation). rRNA pairs were classified as ‘unlinked’ if the distance between each gene was greater than 1500bp, ‘linked’ if the distance was less than or equal to 1500bp. We separated genomes that had a 16S or 23S rRNA gene that started or ended within 1500bp of the beginning or end of its genome and classified these 226 genomes independently to account for the circular nature of bacterial and archaeal genomes. For this subset of genomes, we iteratively adjusted the start and end position of those “edge-case” rRNA genes with respect to genome size and selected the smallest distance between the 16S and 23S rRNA genes as the true distance, using the same formula presented in Supplemental Figure 1. Each genome was classified as ‘unlinked’, ‘linked’, or ‘mixed’ depending on the status of their rRNA genes with ‘mixed’ genomes having multiple rRNA copies with a combination of linked and unlinked rRNA genes. We re-assigned taxonomy to each genome using the SILVA 132 SSU database (clustered at 99%) to maintain a consistent taxonomy between our two datasets. All analyses were done in R version 3.5.1 (R Core Team, 2018). Information on all genomes included in these analyses (including classification of rRNA genes) is available in Supplemental Dataset S1.

### Long-read metagenomic analyses

To investigate the prevalence of unlinked rRNA genes among those Bacteria and Archaea found in environmental samples (including many taxa for which genomes are not yet available), we analyzed long-read metagenomic datasets generated from soil, sediment, activated sludge, anaerobic digesters, and human gut samples. These metagenomic datasets were generated using either the Oxford Nanopore MinION/PromethION (6 samples) or the Illumina synthetic long-read sequencing technology (also known as Moleculo, first described in (Kuleshov et al., 2014), 9 samples). The Moleculo sequences originated from four previously published studies covering: the human gut (Kuleshov et al., 2016), prairie soil (White et al., 2016), sediment (Sharon et al., 2015), and grassland soils (MG-RAST project mgp14596, Flynn et al., 2017). The Nanopore sequences originated from four unpublished studies that studied anaerobic digesters, activated sludge, sediment, and lawn soil. For these samples, DNA was extracted using DNeasy PowerSoil Kits (Qiagen, DE) and libraries were prepared for sequencing using the LSK108 kit (Oxford Nanopore Technologies, UK) following the manufacturers protocol. The libraries were sequenced on either the MinION or the PromethION sequencing platforms (Oxford Nanopore Technologies, UK). Base calling was conducted using Albacore v. 2.1.10 for the lawn soil sample (VCsoil) and Albacore v. 2.3.1 for all other samples (Oxford Nanopore Technologies, UK). Across these 15 samples, we compiled 16,870,533 Nanopore sequences and 846,437 Moleculo sequences with a minimum read length of 1000bp.

We trimmed the first 250bp of each Nanopore sequence to remove low quality regions, but performed no other quality filtering as not all samples included information on sequence quality (some sequences were fasta format). Instead, we relied on our downstream filtering steps to remove sequences of poor quality. Metaxa2 version 2.1 (Bengtsson-Palme et al., 2015) was run on all sequences with default settings to search for SSU (16S rRNA) and LSU (23S rRNA) gene fragments. Taxonomy was assigned to the partial rRNA sequences using the RDP classifier (Wang et al., 2007) and the SILVA 132 SSU and LSU databases (both clustered at 99% sequence identity, Quast et al., 2012). If a sequence contained both 16S and 23S rRNA genes we used the taxonomy with the highest resolution (if the 16S was annotated to family level while the 23S was genus level, we used the 23S taxonomy for both rRNA). Details on each sample, including number of reads and median read lengths, are available in Supplemental Table S1.

We next used a number of criteria to filter the reads included in downstream analyses and to identify taxa with unlinked rRNA genes. We only included those reads in our final dataset that met the following criteria:

1. Included at least 2 domains of the 16S or 23S rRNA genes (Metaxa2 uses multiple HMM profiles targeting conserved regions of the 16S and 23S rRNA genes, each of these regions is referred to as a domain),
2. Included either the last two domains of the 16S rRNA gene (V8|V9) or the first two domains of the 23S rRNA gene (C01|C02),
3. Were ≤4000bp if a 16S rRNA gene and ≤ 6800bp if a 23S rRNA gene (these limits were chosen to accommodate insertions within rRNA genes such as those that occur in Candidate Phyla Radiation (CPR) taxa (Brown et al., 2015), *Nostoc, Salmonella*, and others (Pei et al., 2009)),
4. Could be classified to at least the phylum level of taxonomic resolution.

Of the subset of reads that met these criteria (112 - 878 per Moleculo sample, 3817 - 28056 per Nanopore sample, see Supplemental Table S1 for details), we classified reads as containing unlinked rRNA genes if there was >1500bp between the 16S and 23S rRNA genes, or if there was no 23S domain found 1500bp after the end of the 16S rRNA. We note that, unlike the NCBI gene ranges, Metaxa2 takes strand information into account and translates start and stop locations into sense orientation for SSU and LSU. For our final analyses, we removed reads that could not be classified as linked or unlinked rRNA genes (for instance a sequence with only 300bp after the 3’ end of the 16S rRNA gene) and included only reads that contained a 16S rRNA gene to avoid potentially double counting organisms with unlinked 16S and 23S rRNA genes. All analyses were done in R version 3.5.1 (R Core Team, 2018). Information on all long-read sequences included in these analyses (including classification of rRNA genes) is available in Supplemental Dataset S2.

### Phylogenetic tree combining long-read and NCBI datasets

A phylogenetic tree was created from full-length 16S rRNA sequences by combining both the NCBI complete genomes and representatives of the long-read metagenomic datasets. For the NCBI genome sequences, we selected one 16S rRNA gene sequence per unique species. For the long-read datasets, we first matched the partial 16S rRNA genes recovered by metaxa2 (Bengtsson-Palme et al., 2015) to full-length 16S rRNA gene sequences in the SILVA 132 SSU database (Quast et al., 2012) using the usearch10 version 10.0.240 command usearch_global (settings: -id 0.95 -strand both -maxaccepts 0 -maxrejects 0; Edgar, 2010). The full-length SILVA 16S rRNA genes sequences that matched to the long-read sequences ≥95% percent identity and ≥500bp alignment length were added to the complete genome sequences as representatives of their long-read sequence match. We used 95% percent identity as our cutoff as we found unlinked rRNA gene status to generally be conserved within genera (see below and Supplemental Figure S2). The NCBI and SILVA sequences were then aligned with PyNAST version 0.1 (Caporaso et al., 2010) and the phylogenetic tree was constructed using FastTree version 2.1.10 SSE3 (Price et al., 2009), and plotted with iTOL (Letunic and Bork, 2016).

### Genomic attributes associated with unlinked rRNA genes

All tests for genomic attributes were done with a subset of our complete genome dataset - we reduced the dataset to include only one representative genome per unique species and operon status. For example, if a species had 24 genomes with linked rRNA genes and 3 genomes with unlinked rRNA genes, we retained two genomes total, one linked and one unlinked. Species with heterogeneous rRNA gene status accounted for only 0.71% of species and we found that the presence of unlinked rRNA genes was strongly conserved at the species and genus level (Supplemental Figure S2).

With this set of reduced genomes (3967 genomes in total), we first calculated Pagel’s lambda (Pagel, 1999) to determine whether there was a phylogenetic signal associated with unlinked rRNA genes using the phylosig function of the phytools package version 0.6.60 (Revell, 2011). The results of this test indicated there was a strong phylogenetic signal (lambda = 0.96, p < 0.0001), so we controlled for phylogeny in all of our subsequent tests by using a Phylogenetic Generalized Linear Model for continuous variables (with the function phyloglm in the phylolm package version 2.6, Tung Ho and Ané, 2014).

To determine if taxa with unlinked rRNA genes have a lower predicted growth rate, we calculated the codon usage proxy ΔENC’ (Novembre, 2002; Rocha, 2004), which provides an estimate of minimum generation times (Vieira-Silva and Rocha, 2009). We calculated ΔENC’ with the program ENCprime (Novembre, 2002) with default options, on both the concatenated ORF sequences and concatenated ribosomal protein sequences for each genome following Vieira-Silva and Rocha (2009). To determine if RNaseIII was present in each genome, we used HMMER version 3.1b2 (Eddy, 2011) to search for three RNaseIII pfams (bacterial PF00636, PF14622, and archaeal PF11469) in the translated protein files of each genome. We used the GA gathering cutoffs profile associated with each of these pfams to set all thresholding (--cut_ga).

## Results

### Unlinked rRNA genes occur frequently in complete genomes

We used a set of 12222 “complete” bacterial and archaeal genomes extracted from NCBI in January 2019 to determine how frequently unlinked 16S and 23S rRNA genes occur. We analyzed the distribution of distances between the closest edges of the closest pairs of 16S and 23S rRNA genes (known as the Internally Transcribed Spacer - ITS) in each genome and found that the vast majority of 16S and 23S rRNA gene pairs (98.7%) had an ITS ≤ 1500bp with an average ITS length of 418.7bp (±169.7bp, Figure 2A). However, pairs with ITS lengths > 1500bp showed a scattered distribution of distances, with an average ITS length of 410374bp (±521792bp). Hence, for this classification scheme we called rRNA genes “unlinked” if the ITS was greater than 1500bp. This 1500bp cutoff is in some ways conservative, as the distance between genes in an operon is usually quite low (peaking between −20 and 30bp in most genomes; Moreno-Hagelsieb and Collado-Vides, 2002) and the genes most frequently located between the 16S and 23S rRNA genes encode tRNA, which range from 75 to 90bp in length (Shepherd and Ibba, 2015).

**Figure 2:**
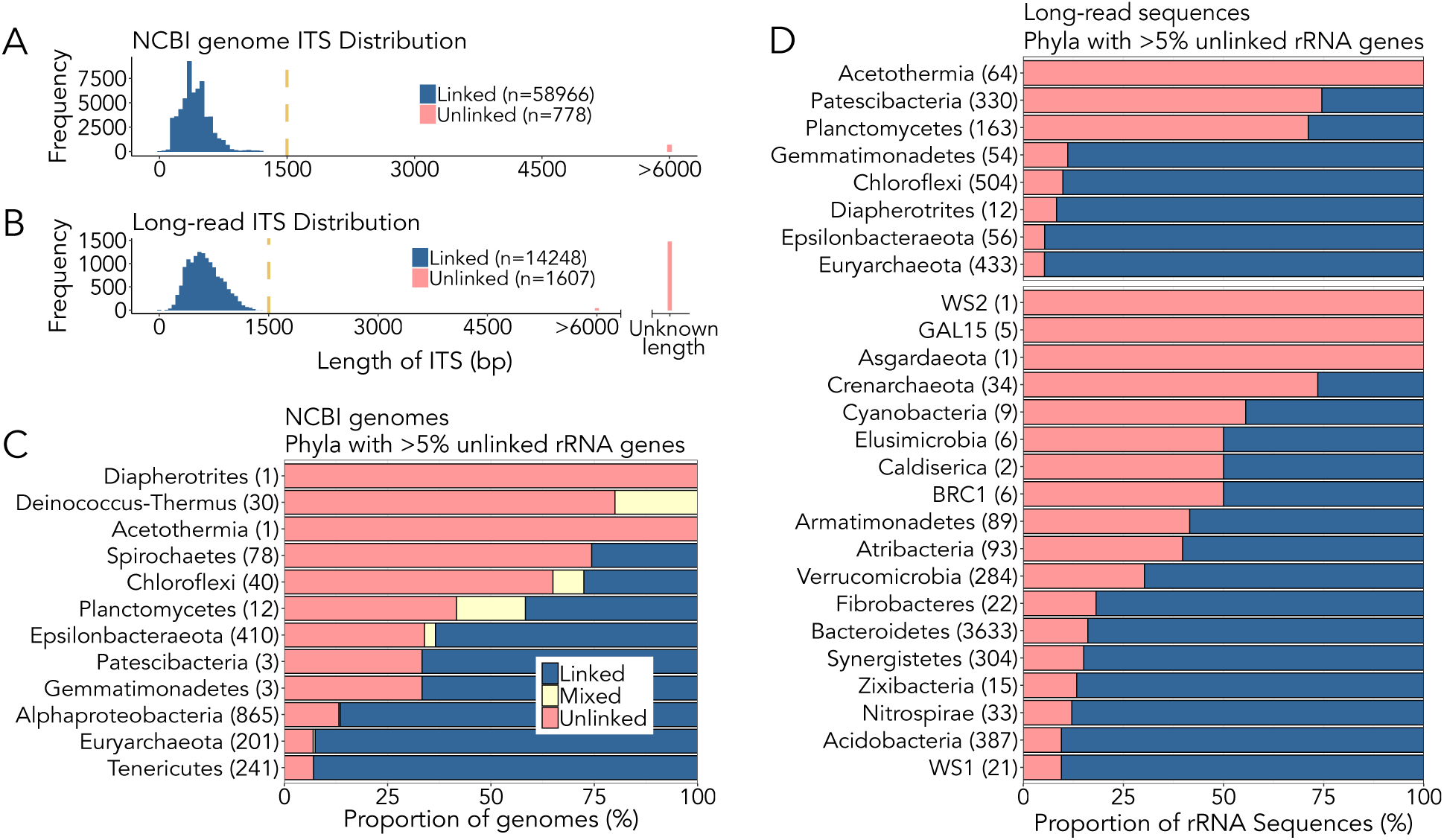
Unlinked rRNA genes can be found in 30 phyla. **A)** Distribution of ITS lengths in complete genomes from NCBI. 98.7% of NCBI rRNA genes have an ITS region ≤ 1500bp in length. The majority of unlinked rRNA genes have an ITS of > 6000bp (682/778) with a mean length of 410374bp (±521792bp). **B)** Distribution of ITS lengths in the long-read sequence dataset. 10.1% of rRNA genes have an ITS > 1500bp. The majority of the unlinked genes have an ITS of unknown length due to sequence length constraints in the long-read dataset (1470/1607). **C)** Within our set of complete genomes from NCBI, 12 phyla had genomes containing at least one set of unlinked rRNA genes in >5% of members. Linked refers to genomes with exclusively linked rRNA genes, unlinked refers to genomes with exclusively unlinked rRNA genes, and mixed refers to genomes with at least one set each linked and unlinked rRNA gene. **D)** By analyzing long-read metagenomic datasets, we confirmed that 8 of the phyla with complete genomes also had unlinked rRNA genes in our environmental or host-associated samples (top portion), and added an additional 18 phyla in which >5% of reads that met our criteria for inclusion in downstream analyses (see Methods) encoded unlinked rRNA genes.

After classifying each rRNA gene pair as linked or unlinked based on the distance between the 16S and 23S rRNA genes, we found that 3.65% of the genomes in our dataset had exclusively unlinked rRNA genes, 0.62% had mixed rRNA gene status (i.e. genomes with multiple rRNA copies that had at least one set of unlinked rRNA genes and at least one canonical, linked rRNA operon), and 95.73% had exclusively linked operons (these numbers do not match up with the per rRNA gene dataset as each genome has a variable rRNA copy number). We found unlinked genomes to be relatively common (present in ≥5% of members) in taxa characterized as having an obligate intracellular lifestyle within the phyla Spirochaetes (genus *Borrelia*), Epsilonbacteraeota (family Helicobacteraceae), Alphaproteobacteria (order Rickettsiales), and Tenericutes (species *Mycoplasma gallisepticum*). However, we also found high proportions of unlinked rRNA genes in phyla that are generally considered to be free-living, such as Deinococcus-Thermus (families Thermaceae and Deinococcaceae), Chloroflexi (family Dehalococcoidaceae), Planctomycetes (families Phycisphaeraceae and Planctomycetaceae), and Euryarchaeota (class Thermoplasmata). Phyla with at least 5% of genomes having exclusively unlinked rRNA genes are shown in Figure 2C.

### Unlinked rRNA genes are widespread in environmental metagenomic data

While the results from our complete genome dataset demonstrate that unlinked rRNA genes are common in some putatively free-living phyla, databases featuring complete genomes do not capture the full breadth of microbial diversity and are heavily biased towards cultivated organisms relevant to human health (Zhi et al., 2012). Just three phyla (Proteobacteria, Firmicutes, Actinobacteria) accounted for >83% of the genomes in our NCBI dataset - even though recent estimates of bacterial diversity total at least 99 unique phyla (Parks et al., 2018). To investigate the ubiquity of unlinked rRNA genes among those taxa underrepresented in ‘complete’ genome databases, we analyzed long-read metagenomic data from a range of distinct sample types. Focusing exclusively on long-read sequences allowed us to span the 1500bp distance required for classification of rRNA genes without the need for assembly. This is important as the repetitive structure of rRNA genes makes it difficult to assemble a mix of non-identical rRNA genes from the short reads typical of most current metagenomic sequencing projects (Yuan et al., 2015).

From our initial long-read dataset encompassing 15 unique samples (∼890,000 Illumina synthetic long reads (also known as Moleculo) and ∼19 million Nanopore reads, with median read lengths of 8858bp and 5398bp, respectively), only 15855 sequences contained rRNA genes and met the criteria we established for the classification of rRNA genes as linked or unlinked (see Methods). Of these reads, we classified 1607 as unlinked, or 10.1% of the dataset (Figure 2B). These long-read metagenomic analyses showed that unlinked rRNA genes are not equally distributed across environments - we found that up to 41% of the taxa in soil had unlinked rRNA genes, whereas other environments had much lower proportions, most notably the human gut, where all sequenced rRNA genes were linked (Figure 3).

**Figure 3:**
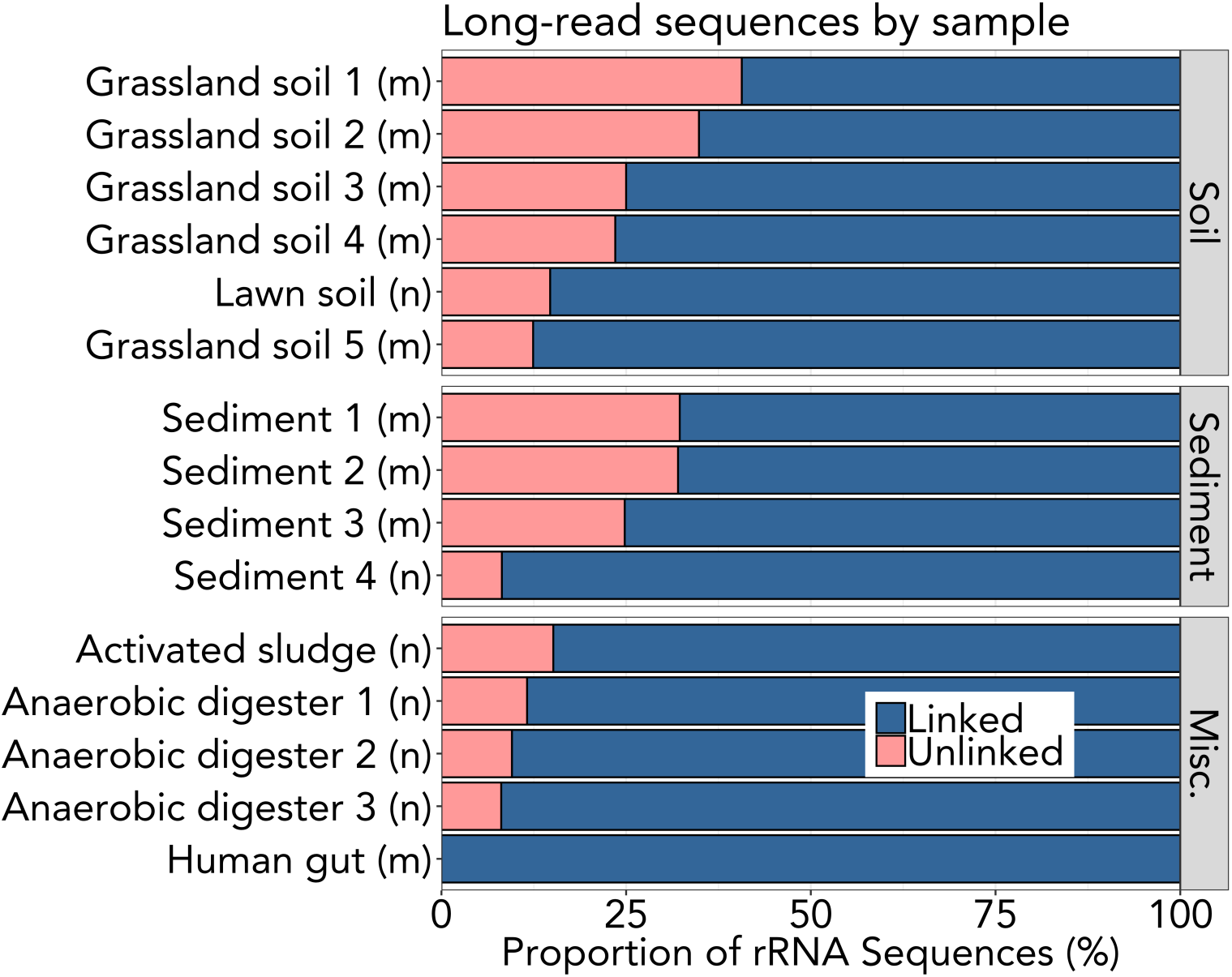
Unlinked rRNA genes have differential frequencies across environments. We found that soils (13-41% unlinked) and sediments (7.7-29%) have more unlinked rRNA genes on average than anaerobic digesters (8.1-8.8%) and the human gut (0%). Results obtained from analyses of Moleculo and Nanopore metagenomic data are indicated with (m) and (n), respectively.

The results from our analyses of the long-read dataset generally mirrored the corresponding results from the complete genome dataset, in that many of the long reads classified as unlinked belonged to the same phyla where unlinked rRNA genes were prevalent in the complete genome dataset (Figure 2). The long-read metagenomic dataset confirmed that members of the phyla Deinococcus-Thermus, Planctomycetes, Chloroflexi, Spirochetes, and Euryarchaeota frequently have unlinked rRNA genes (Figure 2B). The long-read dataset also allowed us to provide additional evidence for unlinked rRNA genes in poorly studied phyla that were represented by only a handful of genomes in our complete genome dataset, such as Acetothermia (1 genome and 64 long-read sequences) and Patescibacteria (3 genomes and 330 long-read sequences).

Using the long-read dataset, we identified 18 additional phyla where unlinked rRNA genes are common, including several candidate phyla (BRC1, GAL15, WS1, WS2) and members of the Candidate Phyla Radiation (Patescibacteria, Figure 2). We also found several clades with high proportions of unlinked rRNA genes that had no representation in our complete genome dataset, including Rikenellaceae RC9 gut group (334/624), Verrucomicrobia genus *Candidatus Udaeobacter* (80/80), Atribacteria order Caldatribacteriales (37/37), Cyanobacteria order Obscuribacterales (4/4), Acidobacteria Subgroup 2 (27/27), Planctomycetes order MSBL9 (40/40), and Chloroflexi class GIF9 (7/7). Overall, we found that 52% of the phyla covered in our combined datasets (37/71) have at least one representative with unlinked rRNA genes.

### Unlinked rRNA genes are strongly conserved

We found that taxa with unlinked rRNA genes are not randomly distributed across bacterial and archaeal lineages - rather, we observed a strong phylogenetic signal for this trait, which we confirmed by calculating Pagel’s lambda (lambda = 0.96, p > 0.001). To highlight this point, we assembled a phylogenetic tree from full-length 16S rRNA gene sequences representing both the complete genome dataset and the long-read metagenomic dataset. We found clusters of related taxa with exclusively unlinked rRNA genes (Figure 4) including: Euryarchaeota class Thermoplasmata, the vast majority of Deinococcus-Thermus, CPR division Patescibacteria, Verrucomicrobia DA101 group, Chloroflexi class Dehalococcoidia, and Alphaproteobacteria class Rickettsiales.

**Figure 4:**
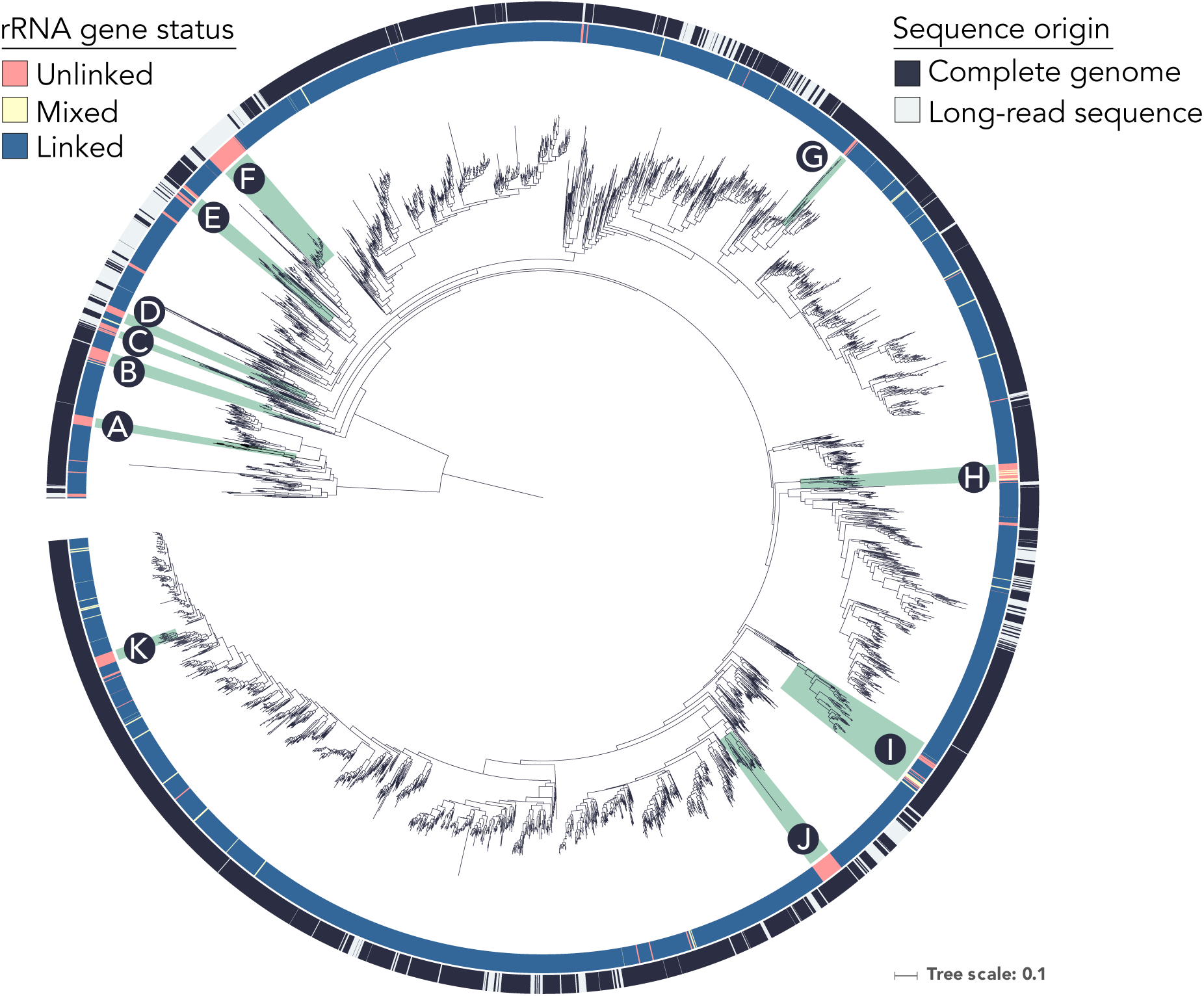
Unlinked rRNA genes occur in coherent phylogenetic clusters. This phylogenetic tree was created from full-length 16S rRNA sequences by combining both the NCBI complete genome and long-read metagenomic datasets (details in Methods). The outer ring indicates which dataset each sequence originated from (complete genomes from NCBI versus long-read sequences from metagenomes), while the inner ring indicates the status of rRNA genes as either linked, mixed, or unlinked. Sequence representatives of the long-read dataset cannot be mixed, as we could not distinguish multi-copy rRNA genes. Clades with high proportions of unlinked members *and* good representation in the tree are indicated in green: A) Euryarchaeota class Thermoplasmata, B) Spirochaetae classes Leptospirae and Spirochaetia, C) Patescibacteria, D) Chlorflexi class Dehalococcoidia, E) Planctomycetes classes Phycisphaerae and Planctomycetacia, F) Verrucomicrobia genus *Candidatus* Udaeobacter, G) Tenericutes genus *Mycoplasma*, H) Deinococcus-Thermus, I) Epsilonbacteraeota genera *Helicobacter* and *Campylobacter*, J) Alphaproteobacteria order Rickettsiales and K) Gammaproteobacteria genus *Buchnera*.

### Genomic attributes associated with unlinked rRNA genes

Given that there are numerous bacterial and archaeal lineages where unlinked rRNA genes are commonly observed, we next sought to determine what other genomic features may be associated with this non-standard rRNA gene arrangement. We treated the presence of unlinked rRNA genes as a binary trait - if a genome had at least one unlinked rRNA gene we counted the genome as “unlinked”. In our NCBI complete genome dataset, we found rRNA gene status to be conserved strongly at the species level - meaning that the majority of species had either exclusively linked or unlinked rRNA genes among their members (Supplemental Figure S2). Therefore, for the following tests, we used a subset of our NCBI complete genome dataset - retaining only a single representative of each species, unless the species had heterogeneous rRNA gene status (0.71% of species), in which case we retained one genome of each rRNA gene status. The analyses were corrected in order to account for the effect of phylogenetic structure in the data (see Methods).

Historically, unlinked rRNA genes have been strongly associated with the reduced genomes of obligate intracellular bacteria, implying that this trait may merely be a side effect of the strong genetic drift and weak selection these taxa experience. To test this hypothesis, we compared the genome sizes of species with linked and unlinked rRNA genes using Phylogenetic Generalized Linear Models (phyloglm). While we found that genomes with unlinked rRNA genes had smaller genomes on average, this difference was not significant (Figure 5, phyloglm p =0.12, means of groups: 4.15Mbp linked, 2.72Mbp unlinked).

**Figure 5:**
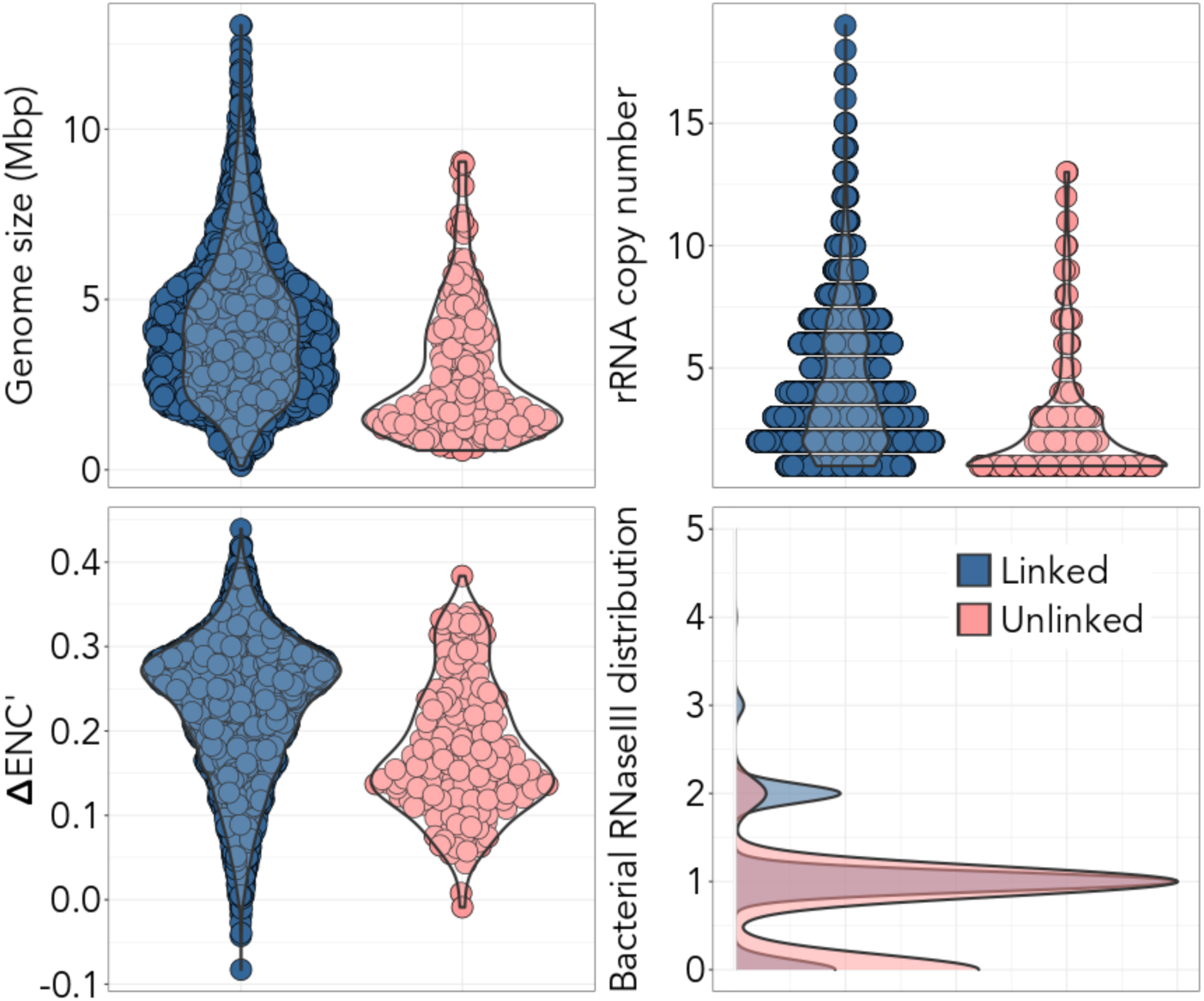
Genomic attributes of NCBI complete genomes based on their rRNA gene status. Linked genomes feature exclusively linked rRNA genes; unlinked genomes have at least one set of unlinked rRNA genes. We calculated these statistics using a subset of our complete genomes with one genome per unique species and rRNA gene status. **A)** Genomes with unlinked rRNA genes have smaller genomes on average, but this difference was not significant (phyloglm p =0.12, means of groups: 4.15Mbp linked, 2.72Mbp unlinked). **B)** On average, genomes with unlinked rRNA genes had significantly fewer rRNA copies (phyloglm p < 0.0001, means of groups: 4.25 copies linked, 2.72 copies unlinked). **C)** Genomes with exclusively unlinked rRNA genes are predicted to have longer average generation times (phyloglm p=0.028, means of groups: 0.23 linked, 0.18 unlinked; as reference E. *coli* has an average ΔENC’ of 0.3). **D)** We found that there were significantly fewer RNaseIII genes in genomes with unlinked rRNA genes (only PF00636 shown, for more detail see Supplemental Figure S3: phyloglm p<0.001, means of groups: 1.0 linked, 0.71 unlinked). This panel is a density plot, which shows the proportional distribution of PF00636 hits for our genome dataset.

The organization of rRNA genes within the same operon facilitates their joint regulation and co-expression at precise stoichiometric ratios. Selection for this trait is expected to be stronger in faster growing Bacteria and Archaea, where, at maximum growth rates, synthesis of the ribosome is the cell’s chief energy expenditure (Gourse et al., 1996). To test this hypothesis, we analyzed the association between the linkage of rRNA genes and traits related to rapid growth in Bacteria and Archaea. On average, genomes with unlinked rRNA genes had significantly fewer rRNA copies (Figure 5, phyloglm p < 0.0001, means of groups: 4.25 copies linked, 2.72 copies unlinked). We also calculated ΔENC’ for each complete genome - a measure of codon usage bias that is negatively correlated with minimum generation time in Bacteria and Archaea (Vieira-Silva and Rocha, 2009). Interestingly, genomes with unlinked rRNA genes were predicted to have significantly longer minimal generation times (Figure 5, phyloglm p=0.028, means of groups: 0.23 linked, 0.18 unlinked). Additionally, in our long-read dataset we found that unlinked rRNA genes were more common in environments typified by slow growth rates; soil and sediment samples had higher proportions of unlinked rRNA genes than samples from anaerobic digesters and the human gut (Figure 3).

RNaseIII separates the precursors of the 16S and 23S rRNA from their common transcript for subsequent maturation and inclusion in the ribosome (Srivastava and Schlessinger, 1990). RNaseIII is not an essential protein in most Bacteria and Archaea and several phyla in which unlinked rRNA genes are common have been reported to not encode RNaseIII (e.g. Deinococcus-Thermus and Euryarchaeota; Durand et al., 2012). Therefore, we checked if there was a significant association between unlinked rRNA genes and the presence of RNaseIII genes. Interestingly, we found that genomes with unlinked rRNA genes were significantly less likely to encode the bacterial form of RNaseIII genes (Figure 5 and Supplemental Figure S3, PF00636: phyloglm p < 0.001, means of groups: 1.0 linked, 0.71 unlinked; PF14622: phyloglm p = 0.007, means of groups: 0.86 linked, 0.66 unlinked). We were unable to check this relationship for archaeal RNaseIII, due to the size of our archaeal dataset (phyloglm failed to converge, only 39 genomes in our dataset had this gene). However, we note that the archaeal RNaseIII PF11469 was found in only two clades that feature exclusively linked rRNA genes (Euryarcheaota family Thermococcaceae and Crenarchaeota family Thermofilaceae).

## Discussion

While unlinked rRNA genes have been documented previously, we have demonstrated that they are far more widespread among Bacteria and Archaea than expected. We found that unlinked rRNA genes consistently occur in 12 phyla using a dataset of complete genomes (Figure 2C), and 18 additional phyla using a dataset of long-read metagenomic sequences obtained from environmental samples (Figure 2D). Interestingly, some phyla were classified as exclusively linked in our complete genome dataset, yet had many members with unlinked rRNA genes in our long-read dataset. For example, while there were no complete genomes in the phylum Verrucomicrobia with unlinked rRNA genes (0/32), 38% of verrucomicrobial rRNA sequences were unlinked (82/217) in our long-read dataset, with the majority of this group closely related to the bacterium *Ca. Udaeobacter copiosus* from the DA101 soil group (Brewer et al., 2016). This highlights the importance of using a combination of complete genomes, where genetic organization and traits can be assessed rigorously, with metagenomic data that allows us to sample the diversity found in selected environments in an unbiased manner. Together, these independent datasets show that unlinked rRNA genes are present across many bacterial and archaeal phyla.

One obvious ramification of the prevalence of unlinked rRNA genes in environmental samples relates to bacterial genotyping using the full rRNA operon. While sequencing from the 16S rRNA gene into the 23S rRNA gene (thus including the ITS region of the rRNA operon) can increase taxonomic resolution and allow strain level identification (Zeng et al., 2012), our work shows that amplicon-based studies dependent on 16S and 23S rRNA genes being located in close proximity may miss a large portion of bacterial and archaeal diversity. We found the average distance between unlinked 16S and 23S rRNA genes in our complete genome dataset to be ∼410kbp, a rather impractical distance to PCR amplify. While strategies which use reads spanning the 16S and 23S rRNA genes to improve taxonomic resolution (e.g. Zeng et al., 2012; Cuscó et al., 2018) are less likely to introduce biases in some environments (e.g. human gut), they will miss many phylogenetic groups in other environments like soil and sediment, where a significant fraction of taxa lack 16S and 23S rRNA genes located in sufficient proximity to be detected with such approaches (Figure 3).

We used our long-read metagenomic dataset to not only bypass the cultivation bias of our complete genome dataset, but to also estimate the abundance of unlinked rRNA genes in a range of microbial community types. Our analyses of the long-read metagenomic dataset show that taxa with unlinked rRNA genes are far more abundant in some environments than others. Most notably, unlinked rRNA genes were far more common in soil (where as many as 41% of rRNA genes detected were unlinked) than the human gut (where no unlinked rRNA genes were detected, Figure 3). The environments with higher proportions of unlinked rRNA genes (soil and sediment) are generally thought to be populated by slower growing taxa (Brown et al., 2016; Vieira-Silva and Rocha, 2009). Likewise, we found that genomes with unlinked rRNA genes have significantly fewer rRNA copies than genomes with exclusively linked rRNA genes, a trait which is correlated with maximum potential growth rate (Vieira-Silva and Rocha, 2009). We also found that genomes with unlinked rRNA genes are predicted to have significantly longer generation times (using codon usage bias in ribosomal proteins as a proxy for maximal growth rates) compared to genomes with exclusively linked rRNA genes. These lines of evidence suggest that unlinked rRNA genes are more common in the genomes of taxa with slower potential growth rates.

The existence of numerous genomes that have unlinked 16S and 23S rRNA genes and the differential frequency of these genomes across environments raise the question of the role and implications of this genetic organization. Upon first consideration, having unlinked 16S and 23S rRNA genes would seem to be disadvantageous given that both rRNA molecules are needed in equal proportions to yield a functioning ribosome. The importance of linkage for identical expression of both rRNA genes should be greater in faster growing taxa, where a higher rate of ribosome synthesis is key to rapid growth and accounts for a large proportion of the cell energy budget (Gourse et al., 1996). Studies in the fast-growing species *E.coli* have shown that, while unbalanced rRNA gene dosage has a slight negative effect on doubling times, balanced synthesis of ribosomal proteins still occurs in most cases (Siehnel and Morgan, 1985). If unequal expression of rRNA subunits is associated with unlinked rRNA genes, it may not confer a selective disadvantage in many environments (like soils and sediments) where longer generation times are the norm, not the exception. For slower-growing taxa, the selection coefficient associated with the effect of linked rRNA genes on growth may be small, because rRNAs are less expressed and rapid growth is a trait under weaker selection. Under these circumstances, unlinked rRNA genes may become fixed in populations by genetic drift. This is more likely to occur in species with small effective population sizes, i.e. few effectively reproducing individuals, where natural selection is not efficient enough to avoid the loss of genes or the degradation of genome organizational traits that are under weak selection (Moran, 2002). This is the most frequent explanation for the occurrence of unlinked 16S and 23S rRNA genes (Rurangirwa et al., 2002; Merhej et al., 2009; Andersson and Andersson, 1999). It fits our observations that many of the taxa we identified with unlinked rRNA genes are restricted to obligate intracellular lifestyles (including members of the phyla Spirochaetes, Epsilonbacteraeota, Alphaproteobacteria, and Tenericutes) or contain signatures of symbiotic lifestyles (CPR phyla; Nelson and Stegen, 2015; Burstein et al., 2016).

However, fixation of mutations due to genetic drift is much less likely to explain the presence of unlinked rRNA genes among the large proportion of free-living taxa that we have identified (including members of the phyla Deinococcus-Thermus, Euryarchaeota, Chloroflexi, Planctomycetes, and Verrucomicrobia). Some of these taxa are abundant and ubiquitous in their respective environments, e.g. the Verrucomicrobia *Ca. U. copiosus* (Brewer et al., 2016) and members of the Rikenellaceae RC9 gut group (Holman et al., 2017). These genomes do not show traits typically associated with genome reduction caused by small effective population sizes, i.e. abundant pseudogenes, transposable elements, or small genomes. Indeed, we found that differences in genome size between species with linked and unlinked rRNA genes were not significant, when accounting for phylogeny. Thus, there is little evidence that the highly conserved trait of unlinked rRNA genes is caused by genetic drift in free-living taxa.

Unlinked rRNA genes could provide a selective advantage in certain circumstances, which could explain their existence in free-living taxa. Transcribing the 16S and 23S rRNA genes separately may eliminate or reduce the need for RNaseIII, which we showed to occur in lower frequencies in taxa with unlinked rRNA genes (Supplemental Figure S3). We also found RNaseIII to be completely absent in the phyla Deinococcus-Thermus and Gemmatimonadetes, both phyla with high proportions of unlinked rRNA genes. Interestingly, there is evidence that the loss of RNaseIII has negative consequences for growth in organisms with unlinked rRNA genes. Recent work has shown that knocking out RNaseIII in *Borrelia burgdorferi* (a spirochete with unlinked rRNA genes) results in a decreased growth rate (Anacker et al., 2018). On the other hand, it is known that some bacteriophages hijack host RNaseIII to process their own mRNA (Gone et al., 2016). In some cases, host RNaseIII can stimulate the translation of infecting phage mRNA by several orders of magnitude (Wilcon et al., 2002), (although other phage appear indifferent to the presence of RNaseIII; Hagen and Young, 1978). Regardless, increased resistance to predation (phage attack) at the cost of reduced maximum potential growth rates is a widely observed ecological trade-off (Bohannan and Lenski, 2000). Finally, recent work has showed that some rRNA loci specialize in the translation of certain types of genes in *Vibrio vulnificus* (Song et al., 2019). It is thus tempting to speculate that unlinked rRNA genes could facilitate the production of heterogeneous ribosomes with a diverse range of characteristics.

## Conclusions

While we do not know why unlinked rRNA genes are so prevalent (especially for those Bacteria and Archaea found in environmental samples for which complete genomes are not yet available), this rearrangement appears to occur more frequently in slower-growing taxa and may be related to the presence of RNaseIII. Regardless, we have shown that 52% of the phyla included in our combined datasets (37/71) have at least one member with unlinked rRNA genes, and that up to 41% of rRNA genes in some environments are unlinked - meaning unlinked rRNA genes are far from atypical anomalies. We have developed hypotheses about potential advantages of unlinked rRNA operons that could be tested experimentally - especially as a number of taxa with unlinked rRNA operons are relatively easy to manipulate in culture (Holland et al., 2006; Devos, 2013).

## Acknowledgements

This research was supported in part by the Chateaubriand Fellowship awarded to T.E.B. from the Office for Science & Technology of the Embassy of France in the United States and a grant to N.F. from the U.S. National Science Foundation (EAR1331828). M.A. was supported by a research grant (15510) from Villum Fonden. A.E. gratefully acknowledges the support of a Leverhulme Trust Research Fellowship (RF-2017-652\2). E.R. was supported by the INCEPTION project (PIA/ANR-16-CONV-0005). We thank Will Trimble for assistance tracking down publicly available Moleculo sequences, Michael Engel for figure design input, and Eric Johnston for introducing the lead author to unlinked rRNA genes.

## Author contributions

TEB, ER, and NF conceived and designed the project and wrote the paper with input from all co-authors. AE, MA, and RK performed the Nanopore sequencing. TEB performed all analyses.

## Data availability

All genomes used in this study were downloaded from NCBI, with assembly IDs listed in Supplemental Dataset S1. All Nanopore data is available at the Sequence Read Archive (SRA) under Bioproject ID PRJNA553237 or the European Nucleotide Archive (ENA) under PRJEB33278. All Moleculo data has been published previously, with publications listed in methods. Classifications and details of both the complete genome and long-read datasets are included in Supplemental Dataset S1 and S2, respectively.

---

**Supplemental Figure S1:**
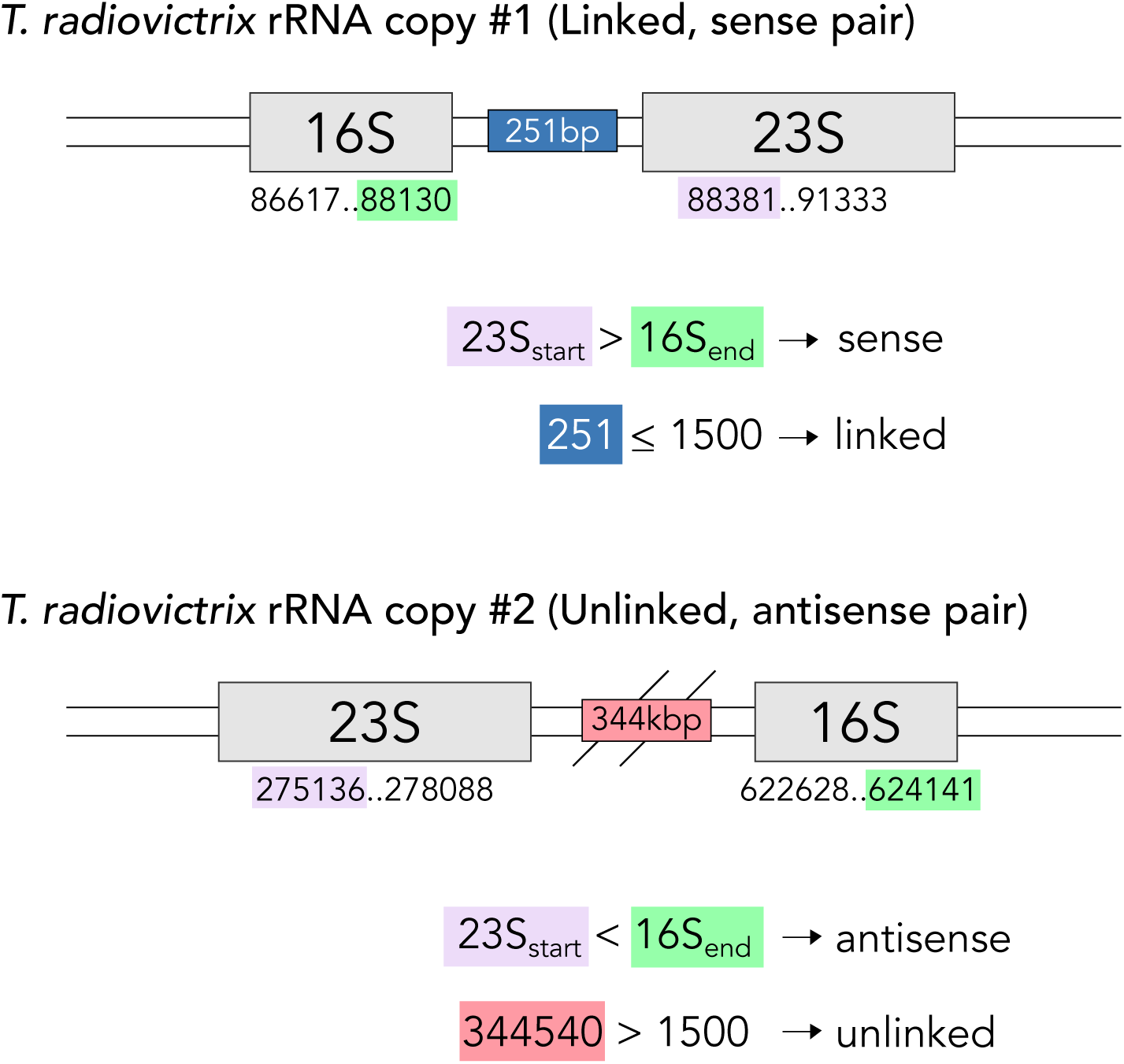
Example of ITS length calculation in *Truepera radiovictrix* DSM17093. To classify the rRNA genes of NCBI genomes, we first used the gene ranges associated with each ORF to pair the 16S and 23S rRNA genes that were closest to each other in each genome. Next, we checked for gene directionality. If the 23S_start_ > 16S_end_ the pair is sense, otherwise antisense. We then calculated the distance between the closest edges of the 16S and 23S (the ITS); if this distance was less than or equal to 1500bp the pair was linked, otherwise unlinked. *Truepera radiovictrix* has an rRNA copy number of two - one pair is linked and sense, the other is unlinked and antisense.

**Supplemental Figure S2:**
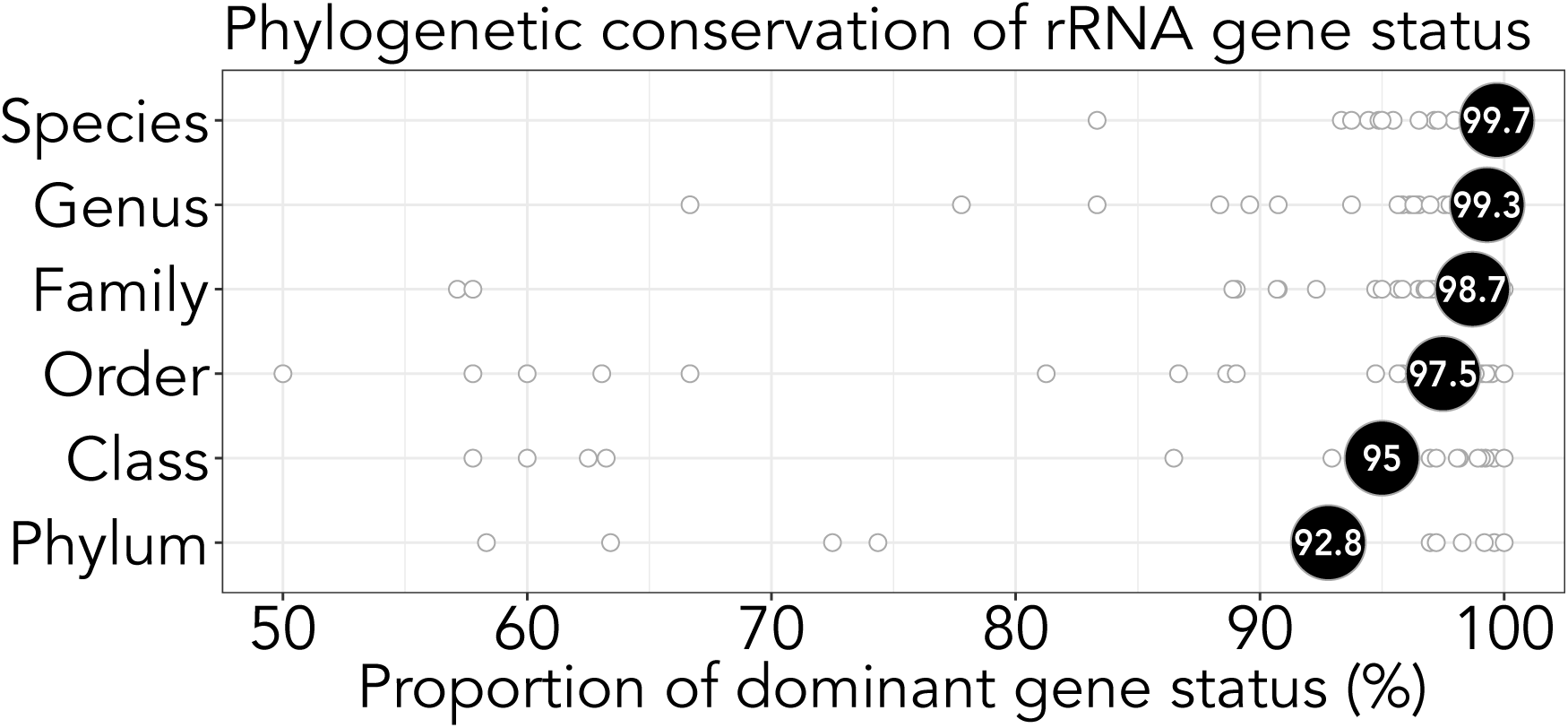
rRNA gene status is phylogenetically conserved, most dramatically at the species and genus levels. Each point represents a unique taxonomic group (for instance, one point at the phylum level corresponds to Chloroflexi). We calculated the proportion of the dominant rRNA gene status for each unique taxonomic group as a measure of trait heterogeneity (for instance, 29/40 Chloroflexi genomes contain an unlinked rRNA gene, making the proportion of the dominant gene status in this phylum 72.5%). The labeled dot corresponds to the average for all taxonomic groups at that specific level. Genera and families with a dominant gene status proportion < 70% are: *Sodalis* (genus), Cellvibrionaceae (family), Spirochaetaceae (family). We also calculated Pagel’s lambda to show that rRNA gene status has a strong phylogenetic signal (lambda = 0.96, p < 0.0001).

**Supplemental Figure S3:**
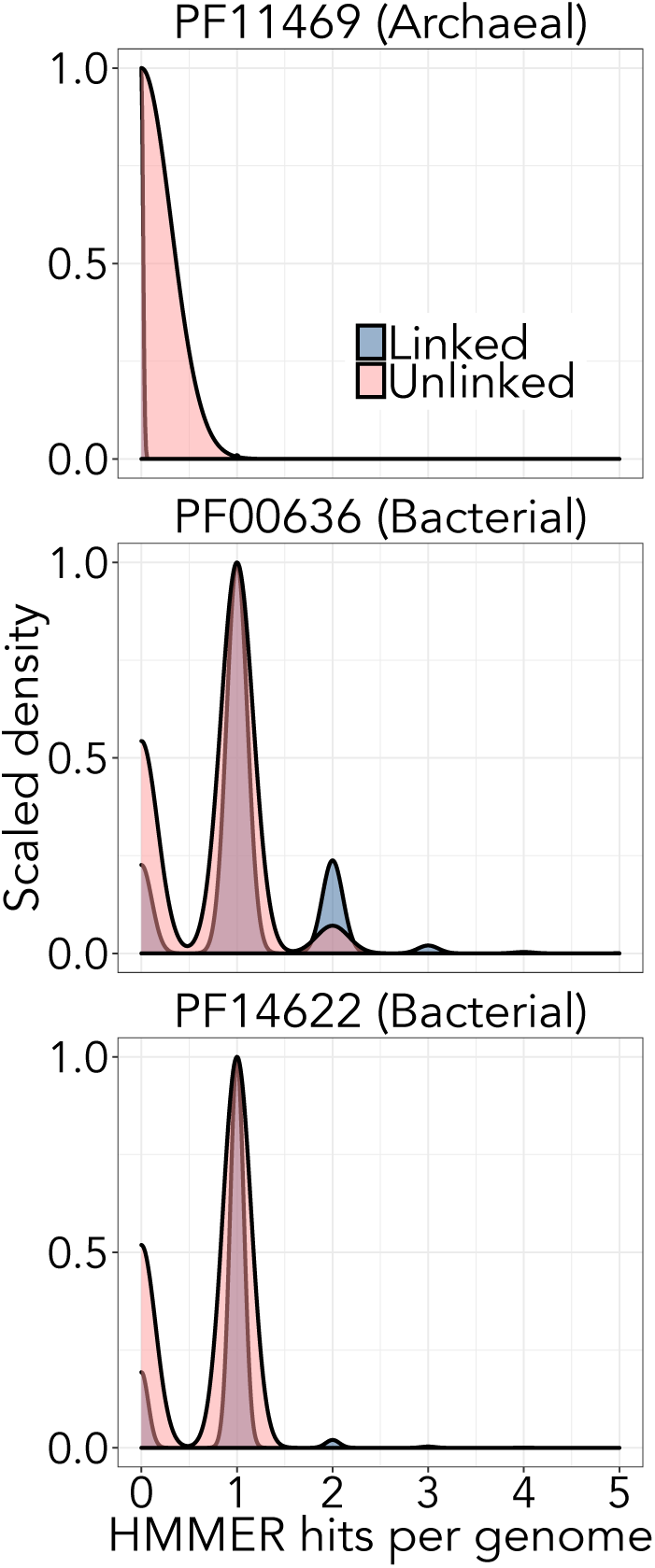
Genomes with unlinked rRNA genes encode fewer bacterial RNaseIII genes. We found that there were significantly fewer bacterial RNaseIII genes in genomes with unlinked rRNA genes (PF00636: phyloglm p<0.001, means of groups: 1.0 linked, 0.71 unlinked; PF14622: phyloglm p = 0.007, means of groups: 0.86 linked, 0.66 unlinked). We were unable to check this relationship for archaeal RNaseIII due to the size of our archaeal dataset. However, the archaeal RNaseIII PF11469 was found in only two clades that featured exclusively linked rRNA genes (Euryarcheaota family Thermococcaceae and Crenarchaeota family Thermofilaceae). We calculated these statistics using a subset of our complete genomes with only one genome per unique species and operon status. These figures are density plots and are scaled to be proportional between the linked and unlinked groups.

